# Comparative plastomics of Amaryllidaceae: Inverted repeat expansion and the degradation of the *ndh* genes in *Strumaria truncata* Jacq

**DOI:** 10.1101/2021.06.21.449227

**Authors:** Kálmán Könyves, Jordan Bilsborrow, Maria D. Christodoulou, Alastair Culham, John David

## Abstract

Amaryllidaceae is a widespread and distinctive plant family contributing both food and ornamental plants. Here we present an initial survey of plastomes across the family and report on both structural rearrangements and gene losses. Most plastomes in the family are of similar gene arrangement and content however some taxa have shown gains in plastome length while in several taxa there is evidence of gene loss. *Strumaria truncata* shows a substantial loss of *ndh* family genes while three other taxa show loss of *cemA*, which has been reported only rarely. Our sparse sampling of the family has detected sufficient variation to suggest further sampling across the family could be a rich source of new information on plastome variation and evolution.

## Introduction

The plastid genome or plastome in land plants is generally conserved in length, structure, and gene content (Wicke et al., 2011). Typical flowering plant plastomes range from 120 to 160 kb, contain 100-120 unique genes, and have a quadripartite structure of two single copy regions (LSC and SSC) separated by two copies of the inverted repeat (IRs) (Jansen & Ruhlman, 2012; Smith & Keeling, 2015). However, exceptions to all of these features have been found. The greatly reduced plastomes of *Pilostyles* range from 11-15 kb and contain only seven functioning genes (Arias-Agudelo et al., 2019). In contrast the plastome of *Pelargonium* × *hortorum* has expanded to 218 kb including unusually long inverted repeats (Chumley et al., 2006). Plastomes deviating from the quadripartite structure have also been reported, either without one copy of the IR (Wojciechowski et al., 2000; Sanderson et al., 2015) or incorporating the entire SSC into the inverted repeats (Sinn et al., 2018).

Rearrangements are often associated with increased repeat content in plastid genomes (Chumley et al., 2006; Haberle et al., 2008; Guisinger et al., 2011). One of the most commonly reported plastome rearrangements is the expansion or contraction of the inverted repeats. Palmer et al. (1985) and Yamada (1991) hypothesized that intramolecular recombination at short inverted repeats located within and around the main IR resulted in the expansion of the *Chlamydomonas reinhardtii* and *Chlorella ellipsoidea* inverted repeats, respectively, while Aii et al. (1997) proposed that recombination at forward repeats within *ycf1* lead to an inversion and the IR expansion in buckwheat (*Fagopyrum* sp.). However, in *Nicotiana*, in the absence of short repeats, Goulding et al. (1996) established two different mechanisms for both short, <100bp, and long, over 10 kb, IR expansions. Short expansions are the result of gene conversion after heteroduplex formation via Holliday junctions. Long expansions are the result of a double-strand break (DSB) in one of the IRs followed by strand invasion and repair against the other IR within the same plastome unit that progresses through the junction incorporating single-copy regions into the inverted repeats. A similar DSB repair process, but via homologous recombination at imperfect, nonallelic repeats between different plastome units, resulted in the reestablishment of the IR in *Medicago* (Choi, Jansen & Ruhlman, 2019). Zhu et al. (2016) attributed the IR expansions in several dicot lineages to the gene conversion mechanism. Furthermore, Wang et al. (2008) proposed that the DSB mechanism of Goulding et al. (1996) can account for shorter expansions as well, such as the development of the monocot-type IR/LSC junction by incorporating *rps19* and *trnH* into the IR.

Various gene losses have occurred during the evolution of the angiosperm plastome (Raubeson & Jansen, 2005; Jansen et al., 2007); a commonly reported example is the loss of *ndh* genes (Song et al., 2017; Silva et al., 2018; Nevill et al., 2019). The loss of *ndh* genes is strongly associated with a change in trophic conditions (Wicke et al., 2013; Graham, Lam & Merckx, 2017); it represents the first step in plastome degradation in heterotrophic plants (Martín & Sabater, 2010; Barrett & Davis, 2012). Apart from plant lineages less reliant on photosynthesis, multiple *ndh* gene losses have also been reported in fully photosynthetic lineages for example in aquatic/semi-aquatic plants (Iles, Smith & Graham, 2013; Peredo, King & Les, 2013; Folk et al., 2020), gymnosperms (Wakasugi et al., 1994; McCoy et al., 2008), Orchidaceae (Kim et al., 2015; Roma et al., 2018), and Cactaceae (Sanderson et al., 2015; Köhler et al., 2020)

Amaryllidaceae J. St.-Hil. (Saint-Hilaire, 1805), is a cosmopolitan family of bulbous geophytes and rhizomatous perennials in Asparagales (Meerow, 2000), comprising approximately 90 genera and over 1,700 species (Meerow, Gardner & Nakamura, 2020) in three subfamilies: Amaryllidoideae, Allioideae, and Agapanthoideae, all of which share an umbellate inflorescence. This family of petaloid monocots contains many horticulturally important genera, including: *Agapanthus, Allium, Amaryllis, Clivia, Galanthus, Hippeastrum, Narcissus, Nerine* (Heywood et al., 2007). Amaryllidoideae, the most diverse subfamily with c. 75 genera (Meerow, Gardner & Nakamura, 2020), has a complex evolutionary history including hybridisation (García et al., 2017; Marques et al., 2017; Meerow, Gardner & Nakamura, 2020) and morphological convergence (Meerow, 2010). Arising from our research in horticulturally important Amaryllidaceae genera (Könyves et al., 2018, 2019a; David & Könyves, 2019) here we report the sequencing and assembly of five Amaryllidoideae species and compare our assemblies with available Amaryllidaceae plastomes from GenBank. We found that most Amaryllidoideae plastomes are of typical organisation, however *Strumaria truncata* has experienced substantial rearrangements and gene losses.

## Materials & Methods

Fresh leaf material was collected from five Amaryllidoideae species at RHS Garden Wisley, UK or from a private collection (Table 1). Herbarium voucher specimens were deposited at WSY. Total genomic DNA was extracted using the QIAGEN DNeasy Plant Mini Kit (QIAGEN, Manchester, UK). Library development and 150bp PE sequencing on an Illumina HiSeq 4000 lane was done by the Oxford Genomics Centre (Oxford, UK). The plastomes were assembled with Fast-Plast v1.2.6 (McKain & Wilson, 2017) and NOVOPlasty v2.7.0 (Dierckxsens, Mardulyn & Smits, 2017). Fast-Plast assemblies were run with a total of 5M, 10M, 20M reads (i.e. 2.5M, 5M, 10M PE reads) and with all available reads. Reads were trimmed to remove NEB-PE adapter sequences. Bowtie reference indices were built with the published *Narcissus poeticus* plastome (MH706763). For the NOVOPlasty assemblies, adapters were trimmed with Trimmomatic v0.36 (Bolger, Lohse & Usadel, 2014) using the same adapter sequences. A *ndhF* sequence of *Na. poeticus* (KT124416) was used as the starting seed and memory was limited to 8 Gb. All other parameters were unchanged. The *Strumaria truncata* NOVOPlasty assembly failed with the *ndhF* seed, a *trnK/matK* sequence of *Na. poeticus* (KC238498) was used instead. Among those assemblies that did not produce consistent results across the different assembly strategies, the large single copy (LSC), the small single copy (SSC), and two inverted repeat (IR) regions were identified in the final Fast-Plast contig and NOVOPlasty assemblies, and the circular plastome was assembled by hand using Geneious v11.1.5 (http://www.geneious.com; (Kearse et al., 2012). The junctions of the inverted repeats and the *ndh* gene sequences in the *S. truncata* plastome assembly were confirmed by Sanger sequencing using the primers and PCR protocols detailed in Table S1 and S2. Coverage analysis of the finished plastomes were done in Fast-Plast. The complete plastomes were annotated by transferring equivalent annotations from the *Na. poeticus* plastome using Geneious v11.1.5, gene and exon boundaries were corrected by hand when necessary.

**Table 1.**
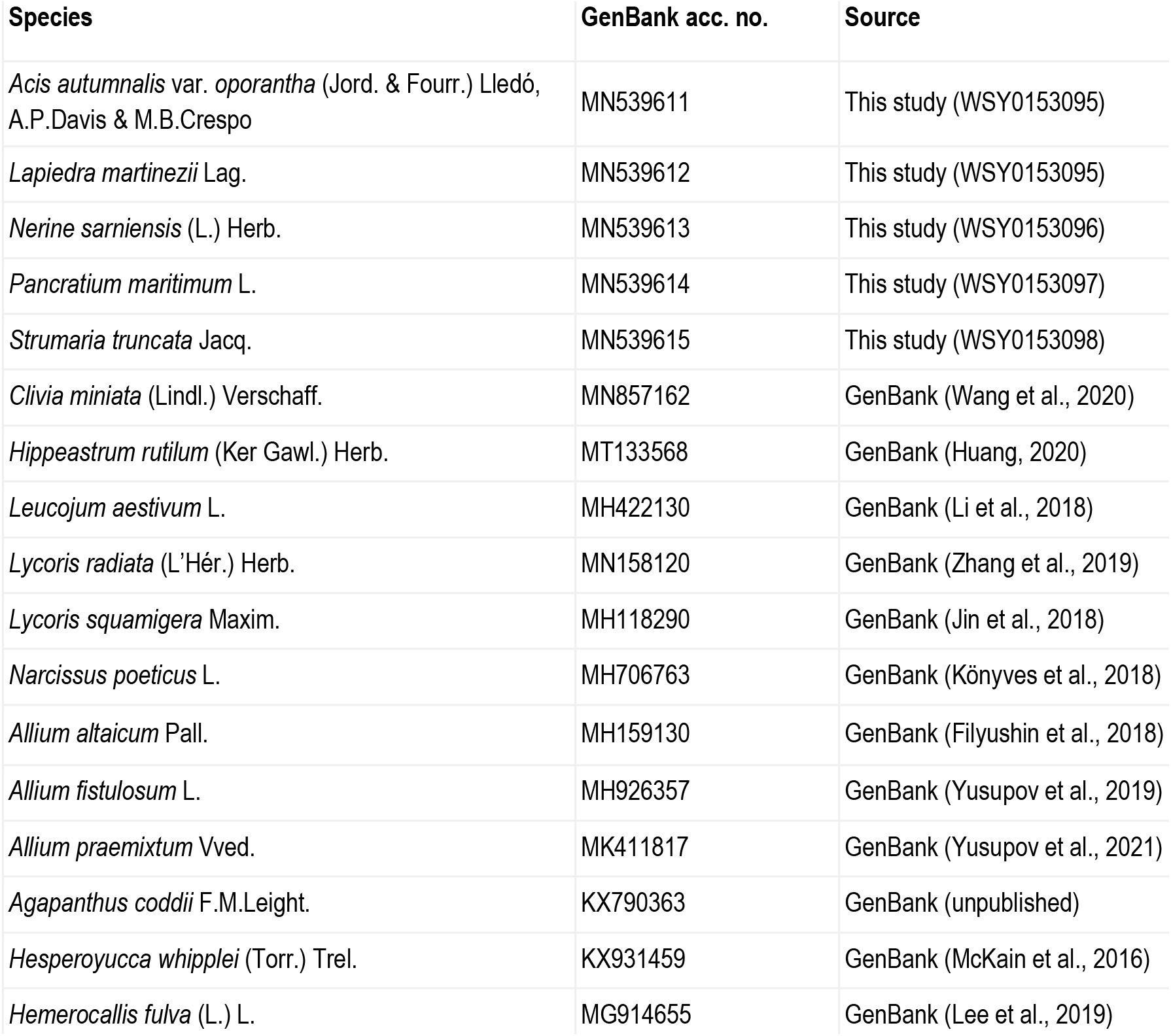
Source information and GenBank accession numbers of the samples included in this study. Herbarium voucher codes (WSY) and publications are listed when available.

The plastomes constructed in this study were combined with ten plastomes representing all three subfamilies of Amaryllidaceae, and a further two Asparagales (Asparagaceae and Asphodelaceae) plastomes from GenBank (Table 1). Sixty-seven coding genes (CDS) that are shared between all samples were extracted from the whole plastomes and aligned with the MUSCLE algorithm v3.8.425 using the default parameters in Geneious Prime 2020.0.5 (https://www.geneious.com). Prior to alignment annotations of GenBank sequences were amended to correct reading frames where necessary (changes listed in Table S3). The alignments of the 67 CDS were concatenated into a matrix of 58230 bp. A maximum likelihood estimate of phylogeny was conducted with RAxML v8.2.11 (Stamatakis, 2014) within Geneious Prime using 1000 bootstrap replicates according to the best-fit model of evolution, GTR+I+G, identified by jModelTest 2 (Guindon & Gascuel, 2003; Darriba et al., 2012).

The inverted repeat boundaries for all samples were identified using the Repeat Finder v1.0.1 plugin in Geneious Prime with default settings. We classified the plastomes into three groups based on the 5’ portion of *ycf1* present in IR_A_, i.e. the structure of the IR_A_-SSC junction (J_SA_), to identify IR expansion events. We did not identify any inversions in the plastomes therefore we tested whether the IR_A_-SSC expansions could have happened through recombination at short inverted repeats present i.e. the mechanism proposed by Palmer et al. (1985) and Yamada (1991) or through short forward repeats which could mediate recombination as mentioned by Choi et al. (2019). For these we searched for short repeats present in both target regions in each species of interest, detailed below, using the Repeat Finder v1.0.1 plugin in Geneious Prime with a minimum of 16 bp length and allowing 10% mismatch. We chose the minimum length and allowed for mismatches as Staub and Maliga (1994) showed evidence of recombination at such repeats in plastids. In *Na. poeticus* and *Pancratium maritimum* (plastome Type B) we screened the region 500-1500 bp from the 5’ end of *ycf1* (the position where J_SA_ in Type A plastomes occurs) and the 500 bp either side of J_SA_ in both species. Repeat Finder did not identify any repeats in *Na. poeticus* so we manually checked the target regions for those homologous with *P. maritimum* to see if this lack is due to our strict search settings. To account for the loss of the *ndh* genes in *S. truncata* we screened for the repeats in *Nerine sarniensis* as well, 500 bp either side of J_SA_ and within *ndhH* where J_SA_ in *S. truncata* is found. The inverted repeat in *S. truncata* contains a 45 bp region downstream of the *ndhH* pseudogene that is absent in other taxa. To identify where this 45 bp region originated we aligned a portion of the *S. truncata* plastome, spanning from *trnN* in IR_B_ to *trnN* in IR_A_, against the same portion of the *Ne. sarniensis* assembly using the MUSCLE algorithm with the default parameters in Geneious Prime. We identified tandem repeats using Phobos v3.3.12 (Mayer, 2006-2010) within Geneious Prime following Joyce et al. (2019) by restricting the search to perfect repeats between 2 and 1000 bp long, with the “remove hidden repeats” setting enabled.

Protein coding genes were categorised as intact, putatively pseudogenized, or lost following Joyce et al. (2019). Briefly, genes were annotated as putative pseudogenes if they contained internal stop codons or the stop codon was missing. Genes were considered lost if less than 30% of the gene was present compared to the other samples. Plastome maps were drawn in OGDRAW v1.3.1 (Greiner et al., 2011).

## Results

Illumina pair-end sequencing for the samples in this study produced 19,015,437-24,547,050 raw paired end reads. All assembly strategies produced consistent assemblies for *P. maritimum*. Fast-Plast set at 20M reads and NOVOplasty assemblies were consistent for *Acis autumnalis* var. *oporantha*. Variation in strategies for assembly for all other plastomes caused differences in length (Table S4), therefore final assemblies were constructed by hand. Sanger sequencing confirmed that the junctions in all five plastomes and the *ndh* genes in *S. truncata* were correctly assembled. Average coverage of the final assemblies ranged from 786× to 1,379× (Table S4). Raw sequence data are available in SRA (BioProject: PRJNA730513); assembled plastomes are available on GenBank (MN539611-15). Amaryllidaceae plastomes (Figure 1) have a quadripartite structure, range from 153,129 to 160,123 bp in length, and contain 70-86 protein coding genes, 38 tRNAs and 8 rRNAs (Table 2).

**Table 2.** Comparison of plastome features in Amaryllidaceae and outgroup samples. Plastomes assembled in this study are in bold.

|  | Amaryllidaceae |  |  |  |  |  |  |  |  |  |  |  |  |  |  | Asparagaceae | Asphodelaceae |
| --- | --- | --- | --- | --- | --- | --- | --- | --- | --- | --- | --- | --- | --- | --- | --- | --- | --- |
|  | Amaryllidoideae |  |  |  |  |  |  |  |  |  |  | Allioideae |  |  | Agapanthoideae |  |  |
|  | ***Acis***<br>***autumnalis***<br>***var.***<br>***oporantha*** | ***Lapiedra***<br>***martinezii*** | ***Nerine***<br>***sarniensis*** | ***Pancratium***<br>***maritimum*** | ***Strumaria***<br>***truncata*** | *Clivia*<br>*miniata* | *Hippeastrum*<br>*rutilum* | *Leucojum*<br>*aestivum* | *Lycoris*<br>*radiata* | *Lycoris*<br>*squamigera* | *Narcissus*<br>*poeticus* | *Allium*<br>*altaicum* | *Allium*<br>*fistulosum* | *Allium*<br>*praemixtum* | *Agapanthus*<br>*coddii* | *Hesperoyucca*<br>*whipplei* | *Hemerocallis*<br>*fulva* |
| GenBank accession number | **MN539611** | **MN539612** | **MN539613** | **MN539614** | **MN539615** | MN857162 | MT133568 | MH422130 | MN158120 | MH118290 | MH706763 | MH159130 | MH926357 | MK411817 | KX790363 | KX931459 | MG914655 |
| Total plastome length (bp) | 157839 | 159022 | 158312 | 160123 | 157566 | 158114 | 158357 | 157241 | 158335 | 158459 | 166099 | 153129 | 153164 | 153226 | 157055 | 157832 | 155855 |
| LSC length (bp) | 85792 | 86756 | 86271 | 86393 | 85648 | 86204 | 86451 | 85657 | 86613 | 86431 | 86445 | 82196 | 82237 | 82162 | 85204 | 86170 | 84607 |
| SSC length (bp) | 18367 | 18540 | 18455 | 16716 | 6904 | 18334 | 18272 | 18180 | 18262 | 18500 | 16434 | 17913 | 17907 | 18042 | 18113 | 18228 | 18508 |
| IR length (bp) | 26840 | 26863 | 26793 | 28507 | 32507 | 26788 | 26817 | 26702 | 26730 | 26764 | 28610 | 26510 | 26510 | 26511 | 26869 | 26717 | 26370 |
| Overall G/C content | 37.7 | 37.8 | 37.8 | 37.8 | 37.8 | 38 | 37.9 | 37.9 | 37.8 | 37.8 | 37.8 | 36.8 | 36.8 | 36.8 | 37.5 | 37.8 | 37.4 |
| Number of tandem repeats | 562 | 605 | 601 | 556 | 591 | 568 | 560 | 567 | 579 | 572 | 543 | 553 | 564 | 561 | 570 | 576 | 666 |
| Sum length of tandem repeats (bp) | 7017 | 7939 | 7922 | 6875 | 7591 | 7024 | 7021 | 7178 | 7140 | 7029 | 6838 | 7066 | 7267 | 7254 | 6975 | 7019 | 8790 |
| Percentage length of tandem repeats | 4.45 | 4.99 | 5.00 | 4.29 | 4.82 | 4.44 | 4.43 | 4.56 | 4.51 | 4.44 | 4.12 | 4.61 | 4.74 | 4.73 | 4.44 | 4.45 | 5.64 |
| N of Intact protein coding genes (unique) | 86 (79) | 86 (79) | 84 (77) | 85 (78) | 79 (70) | 86 (79) | 86 (79) | 86 (79) | 85* (79) | 86 (79) | 85 (78) | 85 (78) | 85 (78) | 85 (78) | 86 (79) | 86 (79) | 86 (79) |
| Pseudo protein coding genes | 0 | 0 | 2 | 1 | 5 | 0 | 0 | 0 | 0 | 0 | 1 | 1 | 1 | 1 | 0 | 0 | 0 |
| Lost protein coding genes | 0 | 0 | 0 | 0 | 4 | 0 | 0 | 0 | 0 | 0 | 0 | 0 | 0 | 0 | 0 | 0 | 0 |
| N of Intact tRNA (unique) | 38 (30) | 38 (30) | 38 (30) | 38 (30) | 38 (30) | 38 (30) | 38 (30) | 38 (30) | 38 (30) | 38 (30) | 38 (30) | 38 (30) | 38 (30) | 38 (30) | 38 (30) | 38 (30) | 38 (30) |
| N of Intact rRNA (unique) | 8 (4) | 8 (4) | 8 (4) | 8 (4) | 8 (4) | 8 (4) | 8 (4) | 8 (4) | 8 (4) | 8 (4) | 8 (4) | 8 (4) | 8 (4) | 8 (4) | 8 (4) | 8 (4) | 8 (4) |

\* *rps19* in IR<sub>A</sub> of *Lycoris radiata* is truncated

**Figure 1.**
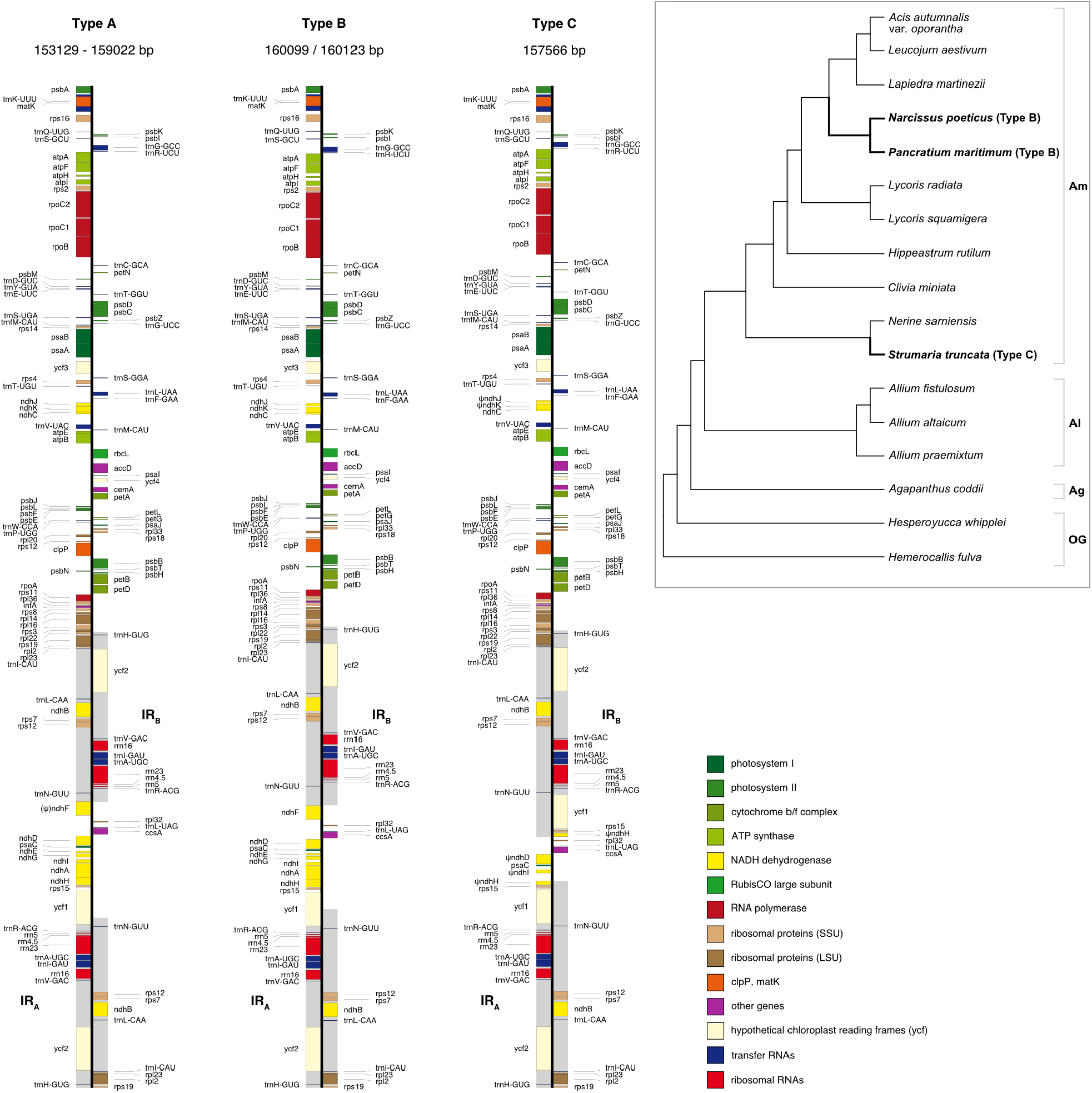
Maps of the three types of plastomes characterised by the 5’ portion of *ycf1* present in IR_A_. Gray shading highlights the IR regions. Genes are coloured according to functional groups shown in the legend. Inset panel shows the relationship between the sampled species based on the RAxML analysis of 67 coding sequences. Type B and C plastomes are highlighted in bold, all other samples had Type A plastomes. Amaryllidaceae subfamilies are indicated on the right: Am=Amaryllidoideae, Al=Alliodideae, Ag=Agapanthodieae, OG=Outgroups.

We identified three different J_SA_ structures (Figure 1) in our samples. Type A was the most frequent junction type found in 14 samples and most likely the ancestral IR junction in Amaryllidaceae as it is shared with both outgroups. The other two J_SA_ structures show two independent IR expansion events. Type B, present in *Na. poeticus* and *P. maritimum*, is a shared IR expansion event, while Type C in *S. truncata* represents a different IR expansion. The 5’ portion of *ycf1* within the IR_A_ ranged from 925-1066 bp for Type A (Figure 2). The Type B plastome had 2649 or 2737 bp of *ycf1* included in IR_A_. The IR expansion in Type C has a different gene arrangement (Figure 2): the entire *ycf1, rps15*, and pseudogenized *ndhH* were included in the IR. The IRs and the SSC ranged from 17853 to 18540 bp and 26370 to 26869 bp, respectively in Type A plastomes. The IRs in the Type B plastomes were longer at 28507 or 28610 bp with shorter SSC of 16434 or 16716 bp. The expanded IRs in *S. truncata* (Type C) were the longest at 32507 bp, while the 6904 bp long SSC was the shortest. The arrangement of the junction between SSC and IR_B_ also showed variation in Type A and Type B plastomes, however this was due to the length variation of the 3’ end of *ndhF* rather than IR expansion/contraction.

**Figure 2.**
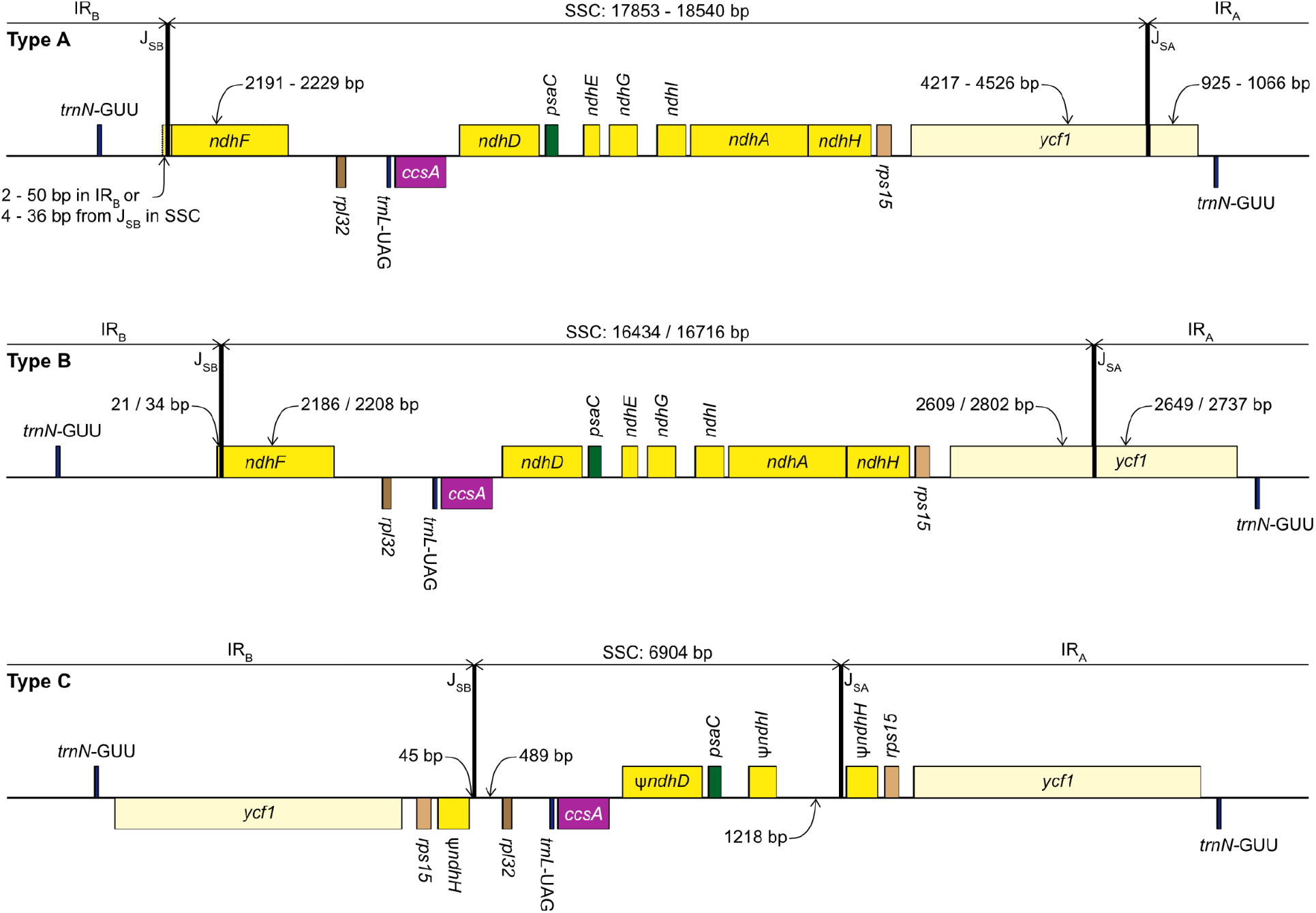
Structure of the junctions (J_SA_, J_SB_) between the inverted repeats (IR_A_, IR_B_) and small single copy (SSC) in the three plastome types identified in this study. The portion of genes included in the IRs and SSC is indicated. Gene order, direction of transcription, and colour code of each gene correspond to Figure 1. ψ indicates pseudogenes.

Repeat Finder identified only one short inverted repeat with our search parameters in *P. maritimum* (Table 3). A similar repeat is present in the *Na. poeticus* plastome, however this has more mismatches than the 10% threshold we set for the search. Both Type B plastomes contain the same short forward repeat, but in *Na. poeticus* this repeat is only partially in our target region.

**Table 3.**
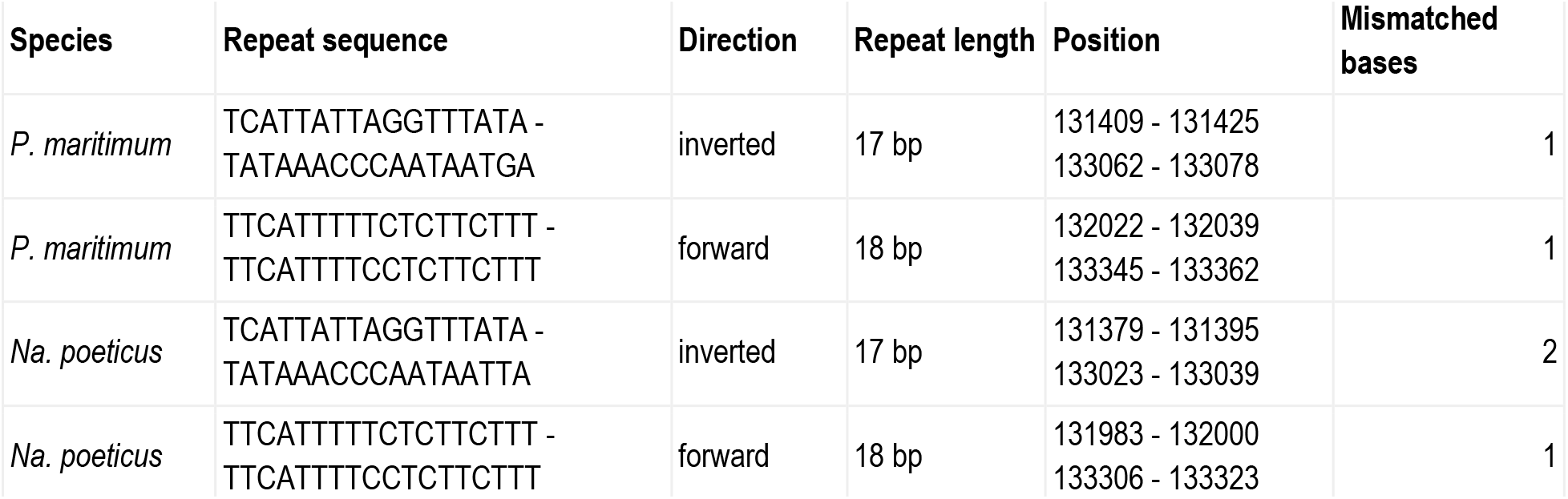
Details of short repeats present at J_SA_ in *Pancratium maritimum* and *Narcissus poeticus*.

The IR expansion in *P. maritimum* and *Na. poeticus* could have happened through recombination of repeats similar to the identified ones in the ancestor of these genera (Figure 3). No short repeats were identified in the *S. truncata* or *Ne. sarniensis* plastomes, therefore the IR expansion in *S. truncata* is likely to be a result of a double-strand break repair as described by Goulding et al. (1996) or Choi et al. (2019). The alignment of the regions between the two copies of *trnN* encompassing the SSC from *Ne. sarniensis* and *S. truncata* showed that the 45 bp present in the *S. truncata* IR and the following single-copy sequence towards *rpl32* is homologous with the *ndhF-rpl32* spacer in *Ne. sarniensis* (Figure S1). These 45 bp represent a second short independent IR expansion in *S. truncata* (Figure 3).

**Figure 3.**
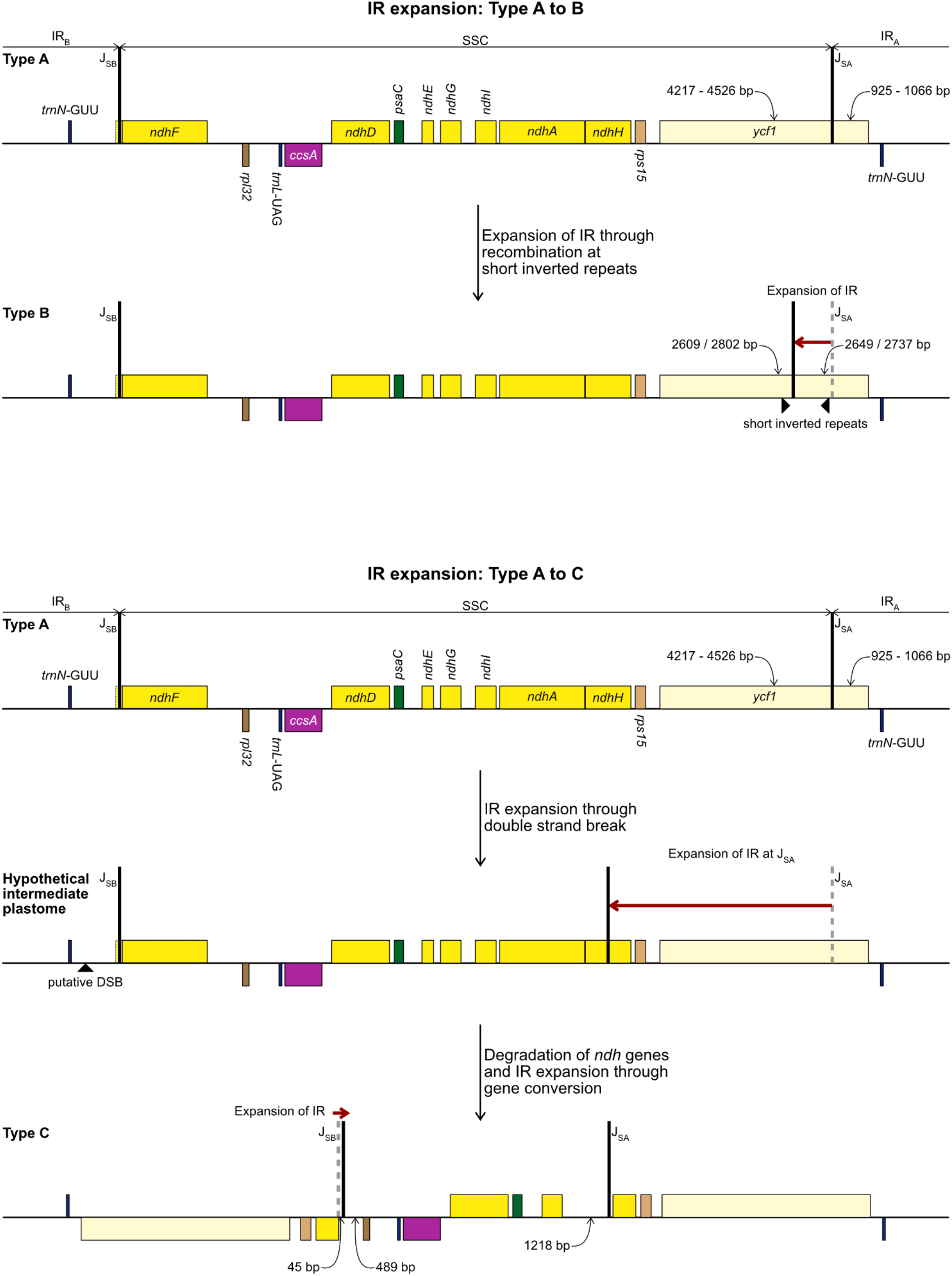
Development of Type B (top) and Type C (bottom) plastome from the ancestral Type A plastome. Steps required to explain the structural changes and the putative mechanisms are detailed in the figure. Gene order, direction of transcription, and colour code of each gene correspond to Figure 1.

The *S. truncata* plastome (Type C) showed a degradation of the *ndh*-suite (Figure 4). Only two *ndh*-genes remained intact, *ndhB* and *ndhC*. The *ndhA, ndhE, ndhF*, and *ndhG* genes were lost, while *ndhD, ndhH, ndhI, ndhJ, ndhK* were pseudogenized. The combined length of the two putative exons of *ndhA* was 21% compared with the other Amaryllidaceae samples. The sequence region was entirely absent for the other genes classified as lost. The *ndhH* gene is 1,182 bp long in Amaryllidaceae. In *S. truncata* this gene is included in the IR as two putative pseudogenes. Each copy is only 592 bp (50%) long from the start codons and the stop codons are missing. The *ndhJ* gene contained an internal stop codon due to transversion mutation (G to T at 49,624 bp), all other putative pseudogenes had frameshift mutations. Furthermore, *ndhF* was classified as a putative pseudogene in *Ne. sarniensis* due to missing stop codon. The *ndhC* gene in *Leucojum aestivum* is missing 47 bp including the start codon at the 5’ end of the gene preceded by 2 N’s potentially indicating scaffolded contigs due to missing data. We did not categorise this gene as a putative pseudogene due to this potential missing data, however we did exclude the gene from the phylogenetic analysis. The *cemA* gene contains frameshift mutations in *Na. poeticus, Ne. sarniensis*, and *P. maritium* due to a homopolymeric A repeat at the 5’ end of the gene. In all *Allium* samples *rps2* is a pseudogene.

**Figure 4.**
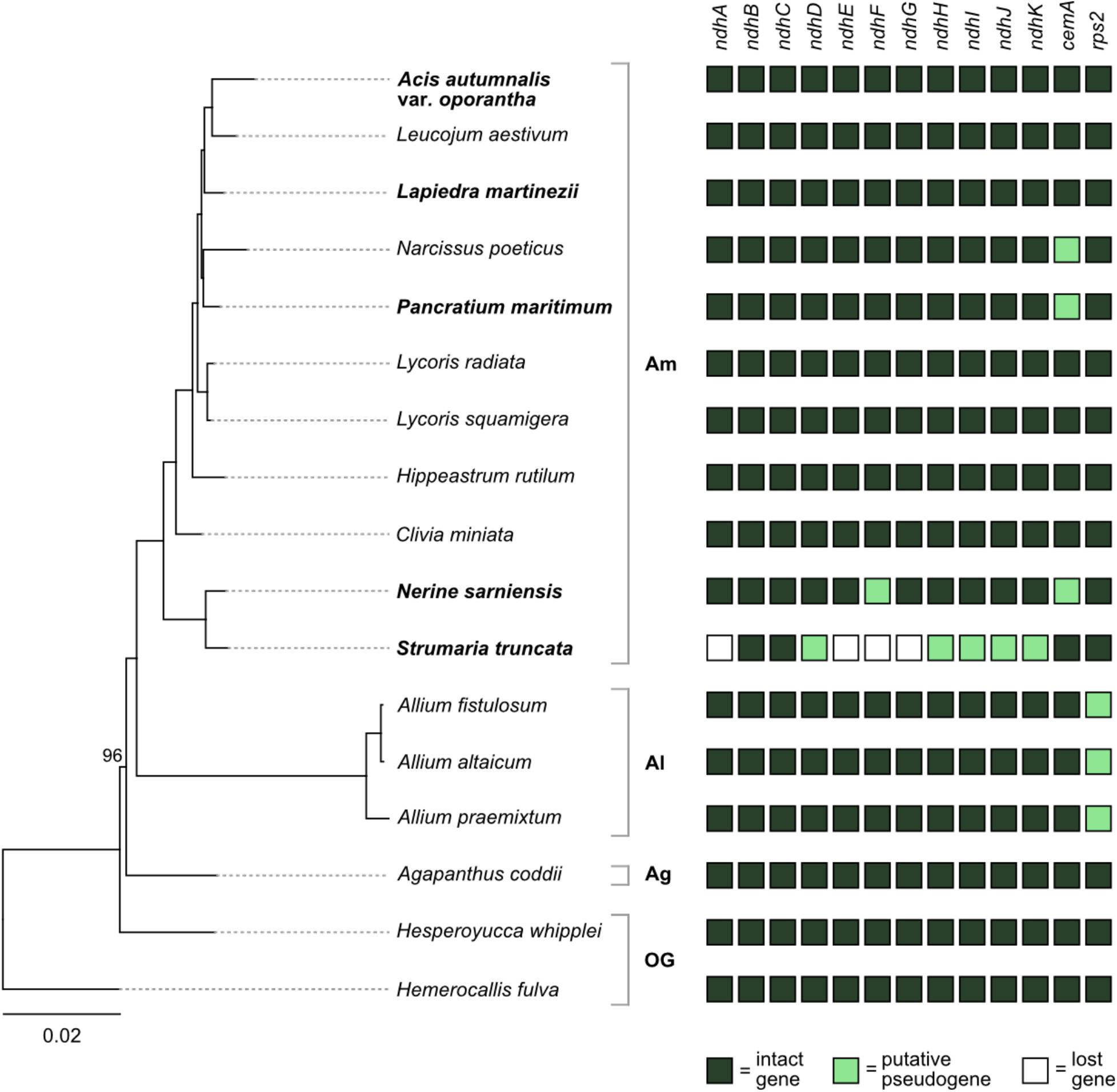
Pattern of gene losses and pseudogenization in Amaryllidaceae and the two outgroup samples plotted against the RAxML tree based on 67 coding sequences. Plastomes assembled in this study are highlighted in bold. All bootstrap support values are 100% apart from the one shown. Amaryllidaceae subfamilies are indicated on the right: Am=Amaryllidoideae, Al=Alliodideae, Ag=Agapanthodieae, OG=Outgroups.

The number of tandem repeats ranged from 543 to 605 in Amaryllidaceae (Table 2). The tandem repeat numbers and their length as percentage of the total plastome (4.1%-5.6%) are similar to those reported by Sinn et al. (2018) for non-rearranged plastomes (420 −732; 3.3%-6.0%). Tandem repeat numbers in Type A plastomes ranged from 560 to 601 (4.4-5.0%) (Table 2). Type B plastomes had 543 (4.1%), 556 (4.3%) tandem repeats in *Na. poeticus* and *P. maritimum*, respectively, and 591 (4.8%) in *S. truncata* (Type C). Moreover, the highest number of repeats, 666, was found in *Hemerocallis fulva* which shows no IR expansion. These changes indicate that an increased number of tandem repeats did not contribute to the IR expansions in Type B and C plastomes.

The phylogenetic analysis based on 67 protein coding genes recovered Agapanthoideae as sister to Allioideae and Amaryllidoideae with 100% bootstrap support (Figure 1). Within Amaryllidoideae the topology is congruent with Rønsted et al. (2012). Sequence alignment of the 67 protein coding genes used in the phylogenetic analysis and the resulting newick tree is available at: https://github.com/kalmankonyves/Amaryllidaceae_plastomes-Strumaria.

## Discussion

The newly sequenced and assembled plastomes of five species from Amaryllidoideae, exhibited three plastome arrangement types based on the portion of *ycf1* within the inverted repeats. Type A- and B-like plastomes have been published previously e.g. *Allium* (Huo et al., 2019) and *Narcissus* (Könyves et al., 2018) respectively, however the Type C plastome, with the largest IR expansion, represents a novel rearrangement in Amaryllidaceae that has not been reported previously.

In typical angiosperm plastomes ∼1000 bp of *ycf1* is included in the IR (Sun et al., 2017). This is similar to the Type A plastome we recognised in Amaryllidaceae. Based on our data we identify the Type A plastome as the ancestral state in the family, as it is shared with both outgroups (Figure 1). The IR expansions that gave rise to the Type B and Type C plastomes represent independent events. In *Na. poeticus* and *P. maritimum* (Type B) the IR expanded to include a larger portion of *ycf1* (2649/ 2737 bp), while in *S. truncata* (Type C) the whole of *ycf1* along with *rps15* and a pseudogenized *ndhH* are contained within the IR. The expansion or contraction of the inverted repeats have been shown to occur in multiple land plant lineages (Wicke et al., 2011; Jansen & Ruhlman, 2012; Zhu et al., 2016) and can often be specific to a few genera within a family (Guisinger et al., 2011; Dugas et al., 2015; Tian et al., 2018; Thode & Lohmann, 2019). In Asparagales IR expansions have been reported in multiple genera in Orchidaceae incorporating genes from the SSC up to and including *ccsA* (Kim et al., 2015, 2020), and in *Eustrephus latifolius*, Asparagaceae (Kim, Kim & Kim, 2016), where the IR expanded to include *ycf1*.

Rearrangements in the plastomes have been associated with an increased number of repeats (Guisinger et al., 2011; Sinn et al., 2018). There was no correlation between the number of tandem repeats and IR expansion in Amaryllidaceae. However, we identified 17 bp inverted repeats in the vicinity of the IR junctions in both *P. maritimum* and *Na. poeticus*. We propose that the IR expansion in these species happened through recombination of similar short inverted repeats in the common ancestor of these genera (Figure 3). The literature offers no consensus on how long short inverted repeats are. The earliest report by Palmer et al. (1985) for *Chlamydomonas reinhardtii*, identified 100-300 bp repeats in the vicinity of the IR junctions, however Aldrich et al. (1988) found 7 bp inverted repeats in *Petunia* and hypothesized recombinations at these led to the IR expansion. We based our search for short inverted repeats on Staub and Maliga (1994) who showed experimental evidence of recombination at 16 bp imperfect repeats in *Nicotiana tabacum* resulting in extrachromosomal elements.

No short repeats were identified in the *S. truncata* or *Ne. sarniensis* plastomes. Therefore, we hypothesize that the IR expansion in *S. truncata* could be a result of a double-strand break repair as described by Goulding et al. (1996) in *Nicotiana* or homologous recombination between different plastome units could also have produced the expanded IR. Choi et al. (2019) proposed that short non-allelic repeats mediated recombination in *Medicago* resulting in the reestablishment of the inverted repeats. As we did not find any suitable repeats, the recombination instead could have happened at homologous genes as shown by Ruhlman et al. (2017) in *Monsonia*. It is also possible that any site of recombination could have been lost during the degradation of the *ndh* genes. We identified 45 bp downstream of the annotated *ndhH* pseudogene to have originated in the *ndhF-rpl32* spacer. This indicates that the IR-SSC organization of *S. truncata* is the result of two independent IR expansion events (Figure 3). Most likely, first a double-strand break originating in IR_B_ got repaired against the complementary strand of IR_A_ with the copy-repair progressing beyond the IR_A_-SSC junction incorporating *ycf1, rps15*, and 592 bp of *ndhH* into the IR. A second small IR expansion originating in IR_B_ further expanded the IR incorporating 45 bp of the region upstream of *rpl32* through the gene conversion mechanisms of Goulding et al. (1996). Although Goulding et al. (1996) called the mechanism responsible for short IR expansion ‘gene conversion’, their description: “Branch migration of a Holliday junction across an IR/LSC junction results in the formation of heteroduplex before this process stalls. Resolution of heteroduplex may then proceed by sequence correction against either strand…” does not preclude this model from applying to expansions at SC/IR junctions with non-coding sequences. Furthermore, ‘gene conversion’ at non-coding sequences removes the constraint of maintaining functional genes.

In *S. truncata* nine out of the 11 *ndh* genes have been lost or pseudogenized (Fig 4). Loss or pseudogenisation of *ndh* genes has been reported in Asparagales both in mycoheterotrophic and autotrophic orchids (Barrett et al., 2014; Kim et al., 2015, 2020; Roma et al., 2018), and in *Allium paradoxum* (2019). We found a further putative *ndh* pseudogene, *ndhF* in *Ne. sarniensis*, which is missing the stop codon. It is possible that this gene is functional and C-to-U RNA editing creates a stop codon at transcription. The pseudogenes in *S. truncata*, on the other hand, are the result of frameshift mutations which are not known to be edited in plastomes (Haberle et al., 2008). The *ndh* genes encode the NADH dehydrogenase-like complex (NDH-1) which regulate photosynthetic electron transport (Peltier, Aro & Shikanai, 2016; Shikanai, 2016). NDH-1 helps adapt photosynthesis under photooxidative stress conditions (Martín & Sabater, 2010) and in environments with fluctuating light intensity (Yamori & Shikanai, 2016). The loss of the *ndh* complex has been reported in plants growing either in high or low light level environments, for example hot desert (Sanderson et al., 2015), submersed (Peredo, King & Les, 2013), or understorey habitats (Graham, Lam & Merckx, 2017; Omelchenko et al., 2019). Furthermore, under optimal growth conditions *ndh* genes appear dispensable (Burrows et al., 1998; Endo et al., 1999; Rumeau, Peltier & Cournac, 2007), and their function could also be assumed by an alternate nuclear encoded system (Wicke et al., 2011) e.g. the AA-sensitive CEF pathway (Sanderson et al., 2015). In the presence of an alternate system, the degradation of the *ndh* genes can have little consequence on plant fitness and mutations can freely accumulate leading to pseudogenisation and loss. Alternatively, *ndh* genes can be transferred to the nucleus and only lost from the plastome, as Roma et al. (2018) showed in *Ophrys*. Based on our data we can only hypothesize the fate of the *ndh* complex. The loss of *ndh* genes from the *S. truncata* plastome compared with *Ne. sarniensis* could be the result of ecological adaptation, as the two species occur in different habitats: *S. truncata* inhabits the semi-arid Succulent Karoo, while *Ne. sarniensis* occurs in the Fynbos, characterised by a wetter, Mediterranen-type climate (Duncan, Jeppe & Voigt, 2020). It is also possible that the loss of the *ndh* genes is a result of a transfer to the nucleus, or the presence of an alternate system. Further sampling of *Strumaria* and *Nerine* species and investigating the nuclear genome will be necessary to identify potential drivers of gene losses.

There were further pseudogenised genes found in Amaryllidaceae: the chloroplast envelope membrane protein encoding *cemA* is pseudogenized in *Ne. sarniensis, Na. poeticus*, and *P. maritimum* due to a homopolymeric A repeat adjacent to the potential initiation codon. A similar pattern of pseudogenization has been shown in *Lilium* (Do & Kim, 2019) and *Cocos nucifera* (Huang, Matzke & Matzke, 2013). The *cemA* gene is not essential for photosynthesis, however in high light conditions *cemA*-lacking mutants of *Chlamydomonas reinhardtii* showed increased light sensitivity (Rolland et al., 1997). The *rps2* gene is functional in Amaryllidoideae and *Agapanthus coddii* but has pseudogenized due to internal stop codons in *Allium*, which has been reported previously by Filyushin et al. (2019), Omelchenko et al. (2019), and Xie et al. (2019).

Kim et al. (2015) and Kim et al. (2020) showed a correlation between the presence of *ndhF* and the organisation of the IR-SSC junctions in Orchidaceae, and proposed that the loss of *ndhF* leads to destabilization of the IR-SSC junctions. While this holds true to *Najas flexilis*, mentioned by Kim et al. (2015), and could be supported by our results, there is also published evidence of the opposite. Plastomes in Thymelaeaceae have all SSC genes apart *ndhF* and *rpl32* incorporated in the IR in the presence of a full set of 11 *ndh* genes (Könyves et al., 2019b; Lee et al., 2020; Liang, Xie & Yan, 2020), and in *Asarum* the IR expanded to encompass the whole of the SSC without the loss of *ndh* genes (Sinn et al., 2018).

The IR expansion in *S. truncata* is likely the result of two independent events, therefore the order in which gene losses and expansions happened are difficult to unravel. It is perhaps more parsimonious to suggest that the degradation of the *ndh* genes or at least the loss of *ndhF* preceded the second IR expansion in *S. truncata*, otherwise the whole of *ndhF* had to be incorporated into the IR and subsequently lost from both IR_A_ and IR_B_. Further sampling in Amaryllidoideae could help establish the sequence of gene loss and IR expansion and any correlation between these events.

The phylogenetic relationship between the subfamilies recovered in this study: Agapanthoideae sister to an Allioideae, Amaryllidoideae clade is congruent with previously published studies based on plastid DNA data (Fay et al., 2000; Givnish et al., 2005, 2018; Pires et al., 2006; Seberg et al., 2012). Moreover, while our sampling represents only a small fraction of the diversity within Amaryllidoideae, the relationship within the subfamily is also broadly congruent with previous studies (Meerow et al., 2006; Rønsted et al., 2012). The long branch leading to *Allium* in our phylogenetic tree is also present in studies using fewer (one to four) plastid genes (Givnish et al., 2005; Seberg et al., 2012; Chen et al., 2013); the reason behind this is beyond the scope of our paper.

## Conclusions

We are still very much in the discovery stage when it comes to patterns of gene loss and duplication in the plastome. Genes linked to a range of functions are implicated in plastome change however, not surprisingly, most of the genes that vary are linked to photosynthesis at one level or another. The loss of so many genes is *Strumaria truncata* is notable and worthy of further phylogenetic investigation through deeper taxon sampling. At this stage we are not in a position to propose driving mechanisms for the change or whether those observed are to some extent evolutionarily neutral to the growing conditions the plants experience. While the family is characterised by bulbous geophytes, the habitats those plants occupy vary enormously from arid through mesic to seasonally inundated. Likewise the light levels tolerated by different species vary from intense sun to quite deep shade. This paper offers a first look at the kinds of variation in plastomes that might be found across the family and indicates this will be a promising area for more detailed investigation.

## Acknowledgements

We thank the High-Throughput Genomics Group at the Wellcome Trust Centre for Human Genetics (funded by Wellcome Trust grant reference 090532/Z/09/Z) for the generation of sequencing data.

