## Supplementary for "Comparative plastomics of Amaryllidaceae: Inverted repeat expansion and the degradation of the *ndh* genes in *Strumaria truncata* Jacq"

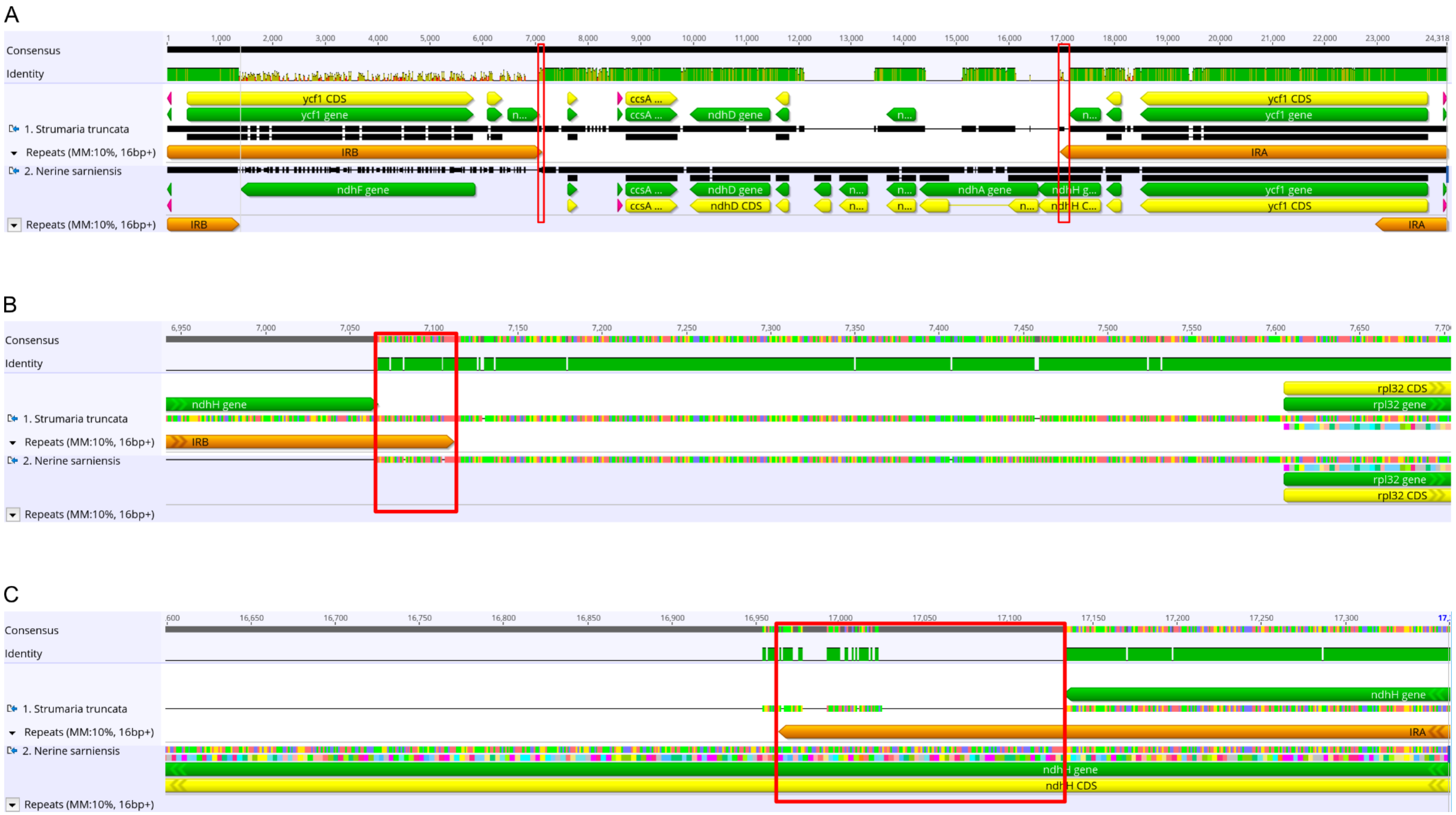


**Figure S1**. Alignment of *Strumaria truncata* and *Nerine sarniensis* plastomes in Geneious Prime 2020.0.5, showing: **(A)** the extracted regions between *trnN* and *trnN* in the IRs, including the complete SSC; **(B)** $J_{SB}$ and **(C)** $J_{SA}$ of *S. truncata.* Red boxes highlight the alignment of the 45 bp sequence region downstream of the *ndhH* pseudogene present in the *S. truncata* IRs. Green bars under 'Consensus' indicate 100% pairwise sequence identity, greeny-brown bars >30% and red bars < 30% pairwise sequence identity. Lack of any bars indicate SNPs and indels in panels B and C. Green arrows indicate gene annotations, yellow arrows show protein coding reading frames, purple arrows show tRNAs, and orange arrows are IR annotations.

**Table S1.** Details of the PCR primers used to amplify and sequence junctions of the inverted repeats and the *ndh* gene sequences in the *Strumaria truncata* plastome assembly. Primers for this study were designed with the Primer3 plugin in Geneious v11.1.5. PCR primers are highlighted in bold, all other primers used as internal sequencing primers.

| Application | Primer | Sequence | Source |
|---|---|---|---|
| Strumaria SSC/IRa | **ndhI_470F** | **CTTTGACCAATGTACCTTGC** | This study |
| | ndhI_rps15_855F | AAATTCATGATAAGAATTCC | This study |
| | ndhI_rps15_1250R | TTAGGAACATCAAGGTACAC | This study |
| | ndhI_rps15_1387F | CTTAATCTTGGAGTAAACAG | This study |
| | ndhI_rps15_1500R | TGATGAGTTAAGCTAGATAG | This study |
| | ndhI_rps15_1723F | ATTATTCGATCCATATCCTG | This study |
| | ndhI_rps15_2334F | TCAGTCTAAGAACACCATGC | This study |
| Strumaria SSC/IRb | **rpl32_120F** | **GGTAGAAACTGACTTTGCTAA** | This study |
| | rps15_rpl32 | AATTGAATAAGTTCATAGTG | This study |
| Strumaria SSC/IRa & IRb | ndhI_rps15_1983R | ATTGTTATATGATCTATTCG | This study |
| | rps15_95R | CRGAGACTYACTTCACATTTGG | This study |
| | **ycf1_end** | **TTTTGAACGAGGAAAAGCAT** | This study |
| Strumaria ndh1 | **ccsA_F** | **ACAAATCAAAGTTTGCAAGGTA** | This study |
| | **psaC_R** | **TAGGATGTACTCAATGTGTACG** | This study |
| | ndhD_F | ATCCATAGAACATCTGACGT | This study |
| | ndhD_R | TTACTGACAGGATTTATCAC | This study |
| Strumaria ndh2 | **ndhJ_F** | **ATGAAATGACAGCAATGGAGTC** | This study |
| | **ndhC_R** | **CCGAATCCGCTATTACATGTTT** | This study |
| Strumaria ndh3 | **psaC_2F** | **CAATAAGATAGACCCATGCTGC** | This study |
| | **rps15_ndhI_R** | **ACTTCTTCCCAACTTGTTTCAC** | This study |
| | ndhI_R | GAATTTGATAAATGTATTGC | This study |
| | psaC_ndhI_F | GTTAGTATTTGTAGTTTCCG | This study |
| LSC/IRa | **rpl2_F** | **CAGAGGGGCTATAATTGGAGAT** | This study |
| | **psbA_R2** | **CGTCCTTGGATTGCTGTTG** | This study |
| | psbA_R1 | CCTTGGTATGGAAGTAATGC | This study |
| | rpl2-psbA-F3 | GGTAARCGYCCYGTAGTAAGAGG | Wang et al. (2008) |
| LSC/IRb | **rps3-F1** | **ATAWATTCYGCAAGAATRTTAGG** | Wang et al. (2008) |
| | **rps3-rpl2-R1** | **AATGGGAAATGCCCTACCTTTG** | Wang et al. (2008) |
| | rps3-F2 | AGTCKGAAACCRAGTGGATTT | Wang et al. (2008) |
| | rps3-rpl2-R2 | GTAGTAAGAGGRGTRGTTATGAACCC | Wang et al. (2008) |

| | | | |
|---|---|---|---|
| LSC/IRb for Nerine and Strumaria | **rps3-F1_G** | **ATAWATTCYGCAAGAATRTTGGG** | Modified from Wang et al. (2008) |
| SSC/IRa & IRb | **trnN** | **CTCTACCACTGAGCTACTGAG** | This study |
| | cp211R_Mod | ACCAAGTTCAACGTTAGCCAGA | Modified from Stoll et al. (2017) |
| SSC/IRa | **cp262_Mod** | **TTTGTATGACCAYCGAGAAACC** | Modified from Stoll et al. (2017) |
| | ycf1_F | GCGATTCCATCGTTTATAATCG | This study |
| SSC/IRb | **ndhF_1318F** | **GGATTAACCGCATTTTATATGTTTC** | Terry et al. (1997) |
| | ndhF_end | GGTCATATAATCGTGGTTAC | This study |
| SSC/IRa poeticus type | **ycf1_888** | **ATACGACATTGATTGACTCTAT** | This study |
| | ycf1_211 | TGGTCGATTCGTGGATAATC | This study |
| SSC/IRb poeticus type | **Pycf1_872** | **GCGTGGTAATTTATACTGATAA** | This study |
| | Pycf1_192 | AGATTCTATTTCCTTTCTGTCTGA | This study |

Stoll A, Harpke D, Schütte C, Stefanczyk N, Brandt R, Blattner FR, Quandt D. 2017. Development of microsatellite markers and assembly of the plastid genome in Cistanthe longiscapa (Montiaceae) based on low-coverage whole genome sequencing. *PloS One* 12:e0178402. DOI: 10.1371/journal.pone.0178402.

Terry R, Brown G, Olmstead R. 1997. Examination of subfamilial phylogeny in Bromeliaceae using comparative sequencing of the plastid locus ndhF. *American Journal of Botany* 84:664.

Wang R-J, Cheng C-L, Chang C-C, Wu C-L, Su T-M, Chaw S-M. 2008. Dynamics and evolution of the inverted repeat-large single copy junctions in the chloroplast genomes of monocots. *BMC Evolutionary Biology* 8:36. DOI: 10.1186/1471-2148-8-36.

**Table S2.** Details of PCR cycling conditions for the amplification of the inverted repeat junctions and the ndh gene sequences in the *Strumaria truncata* plastome assembly.

| | Initial denaturation (temp/time) | Denaturation (temp/time) | Annealing (temp/time) | Extension (temp/time) | Final extension (temp/time) | No. of cycles |
|---|---|---|---|---|---|---|
| Strumaria SSC/IR | 94°C/120s | 94°C/60s | 52°C/30s | 72°C/240s | 72°C/7mins | 35 |
| Strumaria ndh1 | 94°C/120s | 94°C/60s | 53°C/30s | 72°C/180s | 72°C/7mins | 30 |
| Strumaria ndh2 | 94°C/120s | 94°C/30s | 55°C/30s | 72°C/90s | 72°C/7mins | 30 |
| Strumaria ndh3 | 94°C/120s | 94°C/30s | 55°C/30s | 72°C/90s | 72°C/7mins | 30 |
| LSC/IRa | 94°C/120s | 94°C/60s | 56°C/30s | 72°C/180s | 72°C/7mins | 30 |
| LSC/IRb | 94°C/120s | 94°C/60s | 48°C/30s | 72°C/180s | 72°C/7mins | 35 |
| SSC/IR | 94°C/120s | 94°C/60s | 60°C/30s | 72°C/180s | 72°C/7mins | 30 |
| SSC/IR poeticus type | 94°C/120s | 94°C/60s | 52°C/30s | 72°C/180s | 72°C/7mins | 30 |

**Table S3.** Details of reading frame amendments for the Amaryllidaceae plastomes from GenBank.

| sample | gene | original coordinates | new coordinates | note |
|---|---|---|---|---|
| MT133568 | *atpI* | 15131-15382 | 15131-15874 | |
| MN857162 | *rpoC1* | 21194-22492 | 21194-22801; 23558-23989 | added missing exon |
| MN857162 | *petB* | 77128-77133; 78038-78679 | 77245-77250; 78038-78679 | |
| MN158120 | *atpI* | 15235-15678 | 15235-15978 | |
| MN158120 | *ycf1* | 30885-30989 | N/A | renamed to *psbM* |
| MH118290 | *atpI* | 15200-15451 | 15200-15943 | |
| MH118290 | *psbM* | 30857-30982 | 30853-30957 | |
| MH159130 | *rps12* | 67887-68000; 137996-138238 | 67887-68000; 137996-138227; 138770-138795 | added missing exon |
| | | 67887-68000; 97088-97330 | 67887-68000; 96531-96556; 97099-97330 | |
| MH159130 | *petD* | 75432-75995 | 74728-74735; 75482-75995 | added missing exon |
| MH159130 | *rpl16* | 79500-79910 | 79500-79910; 80957-80965 | added missing exon |

**Table S4.** Assembly details for the different plastome assembly strategies. '/' indicates that the two NOVOPlasty outputs differed in length.

| | | | | **NOVOPlasty 2.7.0** | **Fast-Plast 1.2.6** | | | |
|---|---|---|---|---|---|---|---|---|
| | | | | | **5M** | **10M** | **20M** | **all** |
| **Species** | **Total PE reads** | **Final assembled length (bp)** | **Coverage of final assembly** | **length (bp)** | **length (bp)** | **length (bp)** | **length (bp)** | **length (bp)** |
| *Acis autumnalis* var. *oporantha* | 19015437 | 157839 | 786 x | 157881/157839 | 159003 | 159021 | 157839 | 159035 |
| *Lapiedra martinezii* | 19971468 | 159022 | 708 x | 159022 | 146095 | 160059 | 160021 | 160059 |
| *Nerine sarniensis* | 21031067 | 158312 | 1379 x | 158312 | 159609 | 160626 | 160526 | 159540 |
| *Pancratium maritimum* | 24547050 | 160123 | 936 x | 160123 | 160123 | 160123 | 160123 | 160123 |
| *Strumaria truncata* | 23946383 | 157566 | 1197 x | 157566 | Longest contig: 180959 | Longest contig: 181201 | Longest contig: 181186 | Longest contig: 181086 |
